# Multivariate genome-wide association study of rapid automatized naming and rapid alternating stimulus in Hispanic and African American youth

**DOI:** 10.1101/202929

**Authors:** Dongnhu T. Truong, Andrew K. Adams, Richard Boada, Jan C. Frijters, Dina Hill, Maureen W. Lovett, Mark E. Mahone, Eric G. Willcutt, Maryanne Wolf, Pediatric, Imaging, Neurocognition, and Genetics Consortium, John C. Defries, Alessandro Gialluisi, Clyde Francks, Simon E. Fisher, Richard K. Olson, Bruce F. Pennington, Shelley D. Smith, Joan Bosson-Heenan, Jeffrey R. Gruen

## Abstract

Reading disability is a complex neurodevelopmental disorder that is characterized by difficulties in reading despite educational opportunity and normal intelligence. Performance on rapid automatized naming (RAN) and rapid alternating stimulus (RAS) tests gives a reliable predictor of reading outcome. These tasks involve the integration of different neural and cognitive processes required in a mature reading brain. Most studies examining the genetic factors that contribute to RAN and RAS performance have focused on pedigree-based analyses in samples of European descent, with limited representation of groups with Hispanic or African ancestry. In the present study, we conducted a multivariate genome-wide association analysis to identify shared genetic factors that contribute to performance across RAN Objects, RAN Letters, and RAS Letters/Numbers in a sample of Hispanic and African American youth (n=1,331). We then tested whether these factors also contribute to variance in reading fluency and word reading. Genome-wide significant, pleiotropic, effects across RAN Objects, RAN Letters, and RAS Letters/Numbers were observed for SNPs located on chromosome 10q23.31 (rs1555839, multivariate association, p=2.23 × 10^−8^), which also showed significant association with reading fluency and word reading performance (p <0.001). Bioinformatic analysis of this region using epigenetic data from the NIH Roadmap Epigenomics Mapping Consortium indicates active transcription of the gene *RNLS* in the brain. Neuroimaging genetic analysis of fourteen cortical regions in an independent sample of typically developing children across multiple ethnicities (n=690) showed that rs1555839 was associated with variation in volume of the right inferior parietal cortex—a region of the brain that processes numerical information and has been implicated in reading disability. This study provides support for a novel locus on chromosome 10q23.31 associated with RAN, RAS, and reading-related performance.

**AUTHOR SUMMARY:** Reading disability has a strong genetic component that is explained by multiple genes and genetic factors. The complex genetic architecture along with diverse cognitive impairments associated with reading disability, poses challenges in identifying novel genes and variants that confer risk. One method to begin parsing genetic and neurobiological mechanisms that contribute to reading disability is to take advantage of the high correlation among reading-related cognitive traits like rapid automatized naming (RAN) and rapid alternating stimulus (RAS) to identify shared genetic factors that contribute to common biological mechanisms. In the present study, we used a multivariate genome-wide analysis approach that identified a region of chromosome 10q23.31 associated with variation in RAN Objects, RAN Letters, and RAS Letters/Numbers performance in a sample of 1,331 Hispanic and African American youth in the Genes, Reading, and Dyslexia (GRaD) Study. Genetic variants in this region were also associated with reading fluency in GRaD, and differences in brain structures implicated in reading disability in a separate sample of 690 children. The gene, *RNLS*, is located within the implicated region of chromosome 10q23.31 and plays a role in breaking down a class of chemical messengers known to affect attention, learning, and memory in the brain. These findings provide a basis to inform our understanding of the biological basis of reading disability.

## INTRODUCTION

Reading disability (RD, also known as developmental dyslexia) is the most common neurodevelopmental disorder diagnosed in school aged children, with a prevalence of 5-17% (1). RD is characterized by life-long difficulties in reading despite normal intelligence and educational opportunity. The etiological basis of RD is complex and attributed to both genetic and environmental factors. The genetic component of RD is strong, with family and twin-based studies estimating moderate to high heritabilities > 0.50 (2, 3). To date, at least nine putative susceptibility loci for RD (DYX1-9) have been identified through linkage mapping and replicated in several independent populations, along with several candidate risk genes including *KIAA0319*, *DCDC2*, and *DYX1C1* (4-6). However, variants in these candidate genes and loci do not account for a substantial portion of the estimated heritability (7). In addition, genome-wide analyses predicated on quantitative measures of reading ability and case-control status for RD have identified novel variants associated with both, but none have attained genome-wide significance (8-10). This could potentially be explained by low sample sizes in the studies, but another reason may be due to the phenotypic heterogeneity of the reading (dis)ability phenotype.

One approach to studying the genetics of complex trait disorders such as RD is to examine relevant endophenotypes. An endophenotype is a quantitative trait measure that is correlated with a disorder or trait of interest due, in part, to shared (pleiotropic) underlying genetic influences. For a phenotype to be considered an endophenotype for a complex trait disorder, it must have a genetic component, be independent of clinical state (affected or unaffected), co-segregate with disorder status in a family, and have reproducible measurements (11-13). Endophenotypes are conceived as being closer to underlying biology than the corresponding complex disorder. This could improve the statistical power to identify genetic variants through larger effect sizes. Two potential endophenotypes for reading proficiency that satisfy the above criteria are rapid automatized naming (RAN) and rapid alternating stimulus (RAS).

RAN tasks require sequential naming of visually presented, familiar items (e.g. objects, letters, numbers, or colors) as quickly and accurately as possible. RAN has been described as a “mini-circuit” of the reading network that taps into cognitive subprocesses critical for reading such as automatic attentional processing (14, 15). Reading and RAN performance are moderately correlated (0.28-0.57) with deficits in RAN performance observed in 60-75% of individuals with RD (16-18). Furthermore, performance on RAN in kindergarten is predictive of later reading fluency and is stable through elementary school (19, 20). RAN deficits even persist into adulthood (21, 22).

RAS tasks are similar in overall design to RAN tasks, but involve sequential naming of items across *alternating* stimulus categories, again as quickly and accurately as possible (i.e. consistently alternating letters and numbers). RAS tasks aim to evaluate ability to direct attention while performing an automatic task like sequential naming (23). Like RAN, RAS performance can differentiate poor from typically developing readers and is highly predictive of later reading ability (23, 24).

Like RD, RAN has a strong genetic component with heritability estimates ranging from 0.46-0.65 (25, 26). To date, several family-based studies have been conducted to identify novel genetic loci linked to RAN and RAS performance, and to determine whether previously identified DYX loci show linkage and/or association with these potential endophenotypes. The DYX8 region, located on chromosome 1p, showed significant linkage with a composite score of RAN Objects and RAN Colors in a sample of 8 extended multiplex families with at least four people affected with dyslexia (27). In a German sample consisting of 1,030 individuals from 246 families with a history of dyslexia, a multipoint, variance component analysis identified a region of chromosome 6p21 in close proximity to (but not within) the DYX2 region showing significant linkage to a composite score of RAN Objects and RAN Colors (28); however, no linkage signals were identified for RAN Letters or RAN Numbers, respectively, in the sample (28). In a separate study of 1,956 individuals in 260 dyslexia families from the United States, strong linkage was observed on chromosome 2p (within the DYX3 locus) for RAN Letters and chromosome 10q for RAN Colors (29). Moderate linkage signals were observed for RAS Letters/Numbers on chromosome 1p (overlapping with DYX8), chromosome 12q for RAN Numbers and RAS Letters/Numbers/Colors, and chromosome 6q for a composite score of RAS Letters/Numbers and Letters/Numbers/Colors (29). In a sib-pair study of a Dutch RD sample consisting of 108 nuclear families, a composite measure of five RAN measures (Pictures, Colors, Numbers, Small Letters, and Capital Letters) showed weak linkage to the DYX3 and DYX8 loci (30). Candidate variant association studies of RAN measures are limited, but nominal effects of variants within the candidate RD gene, KIAA0319, were reported in a case/control dyslexia sample of 180 Dutch children and a separate Chinese sample of 393 children (31, 32). A nominal effect was also observed for a variant located in SLI1, a locus linked to specific language impairment on chromosome 16q (33). Overall, the above studies support RAN and RAS as endophenotypes for RD, showing that genetic studies targeting RAN and RAS tasks can identify both novel and previously implicated genetic loci involved in RD susceptibility.

Genetic studies of RAN, RAS and RD were previously conducted on samples of largely European descent, and it is unclear whether the same genomic regions and variants contribute to the genetic architecture of these traits in other ethnic populations—specifically populations of Hispanic and African descent who are largely underrepresented in the genetic analysis of RD. Furthermore, the majority of studies focused on linkage mapping, while the limited number of association studies on RAN and RAS directly targeted the strongest candidate genes and/or variants identified for RD from the available literature. The present study aims to identify shared genetic factors that contribute to covariance across RAN Objects, RAN Letters, and RAS Letters/Numbers using a multivariate genome-wide association study in 1,331 unrelated children participating in the Genes, Reading, and Dyslexia (GRaD) study—a case/control sample of RD among Hispanic and African American youth. The present study builds on the previous literature that investigated genetic underpinning of RAN and RAS performance as univariate traits. However, a univariate design can miss underlying covariance across two or more correlated traits and therefore has low sensitivity for detecting shared genetic factors (34). We hypothesize that the correlations between RAN and RAS performance can partially be attributed to shared genetic factors (25). A multivariate genetic analysis allows us to leverage covariance across phenotypes and increase statistical power to identify the presence of pleiotropy across different RAN and RAS tasks (34).

## RESULTS

### Multivariate and Univariate Associations for RAN and RAS

Significant Pearson correlations across RAN Objects, RAN Letters, and RAS Letters/Numbers were observed in the GRaD sample ranging from 0.621–0.781 (p <0.001; S1 Table). Multivariate GWAS of RAS Letter/Numbers, RAN Objects, and RAN Letters was then conducted using the R package, MultiPhen, to identify pleiotropic factors that may contribute to the high phenotypic correlations across RAN and RAS tasks. The model corrected for the effects of age, sex, socioeconomic status (SES), and the first 10 principal components (generated from genome-wide SNP data) to correct for population stratification (S2 Table, S1 Figure). Multivariate analysis of RAS Letter/Numbers, RAN Objects, and RAN Letters revealed a genome-wide significant effect for rs1555839 (p = 2.23 × 10^−8^; Figure 1, S2 Figure, Table 1) located approximately 5 kb upstream from Ribosomal Protein L7 Pseudogene 34 (*RPL7P34*), a long non-coding RNA (lncRNA) on chromosome 10 mapping between the genes *LIPF* and *LIPJ*. Additional markers approaching significance were also clustered within a 70 kb region of chromosome 10q23.31 spanning *RPL7P34* and the gene Renalase (*RNLS*) (Table 1). Post-hoc univariate examination of RAN Objects, RAN Letters, and RAS Letters/Numbers was conducted against the top SNPs identified in the multivariate analysis to determine whether a specific variable(s) may be driving the multivariate signal. The analysis showed that the markers located on chromosome 10q23.31 had a statistically significant effect across all RAN and RAS tasks (Table 1). The strongest association in the chromosome 10q23.31 region was with marker rs1555839 (β = −0.32, p = 1.04 × 10^−9^) for RAS Letters/Numbers.

**Figure 1:**
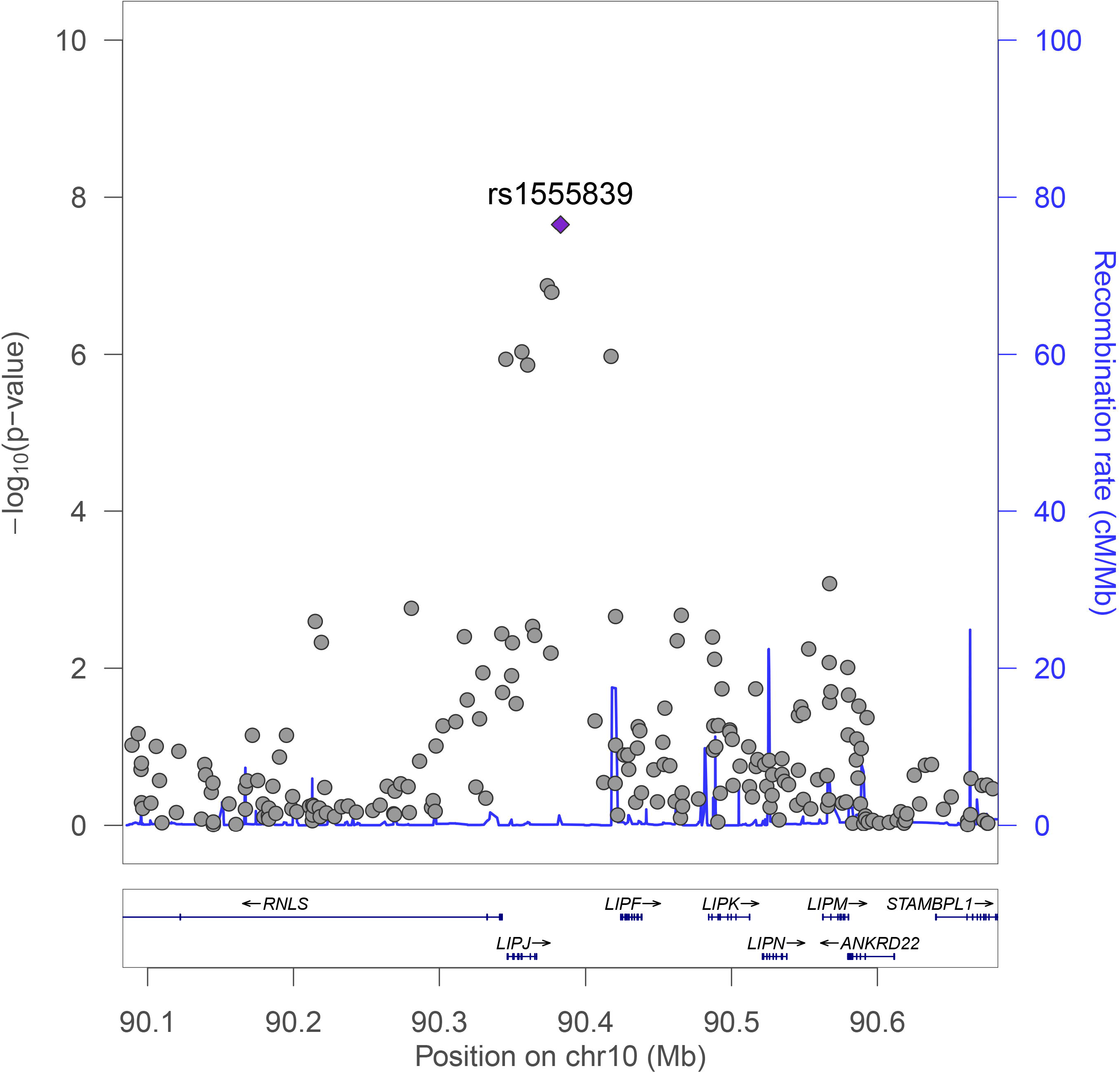
LocusZoom plot of genomic region surrounding genome-wide significant SNP, rs1555839, in multivariate GWAS for RAN Objects, RAN Letters, and RAS Letters/Numbers. LocusZoom plot represents −log10*p* (left y-axis) of each genotyped SNP (gray dots) surrounding the top associated SNP, rs1555839 (purple dot). Recombination rate (right y-axis) is represented by the blue overlay. LncRNA pseudogene, RPL7P34 (chr10: 90,377,980-90,378,691), is not represented in this plot.

**Table 1:**
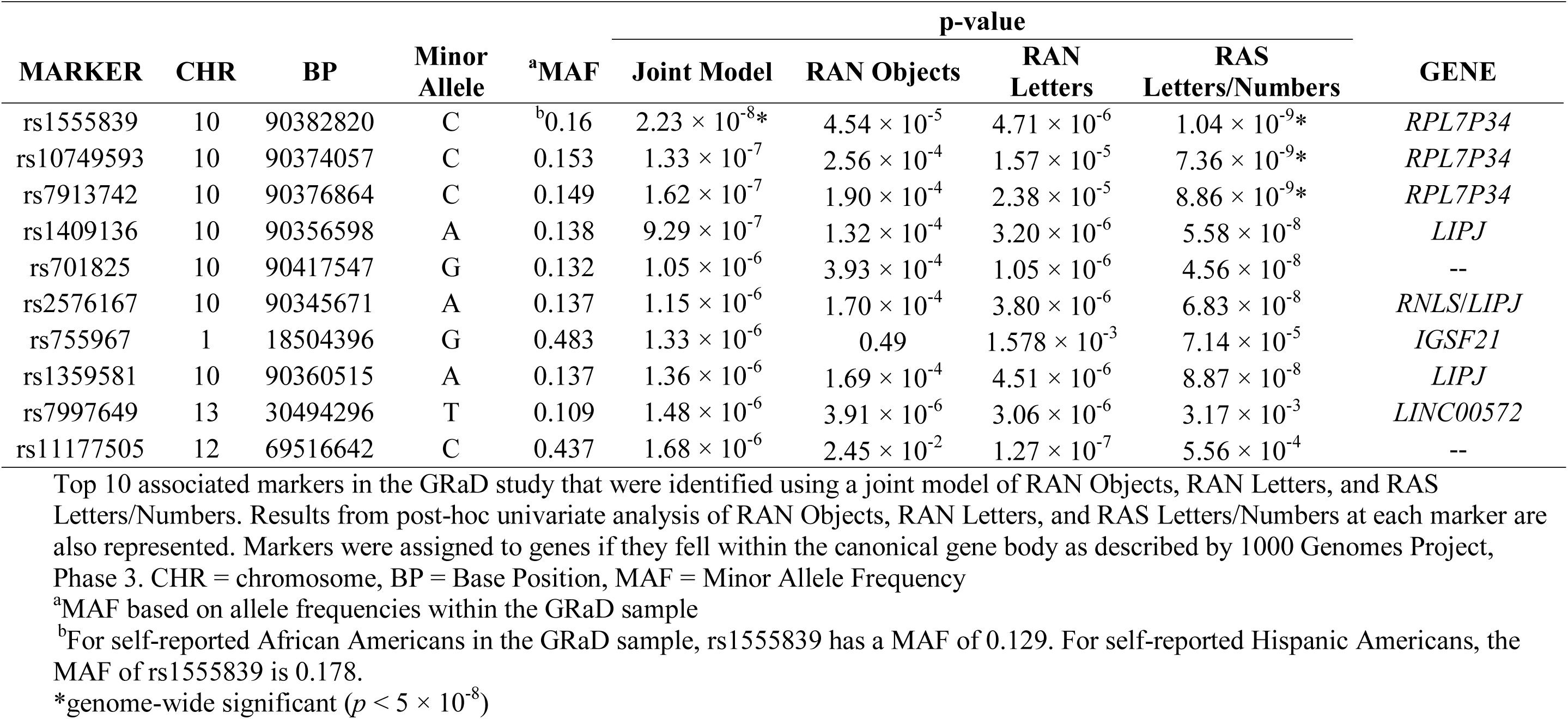
Multivariate GWAS results.

Top 10 associated markers in the GRaD study that were identified using a joint model of RAN Objects, RAN Letters, and RAS Letters/Numbers. Results from post-hoc univariate analysis of RAN Objects, RAN Letters, and RAS Letters/Numbers at each marker are also represented. Markers were assigned to genes if they fell within the canonical gene body as described by 1000 Genomes Project, Phase 3. CHR = chromosome, BP = Base Position, MAF = Minor Allele Frequency

a MAF based on allele frequencies within the GRaD sample

b For self-reported African Americans in the GRaD sample, rs1555839 has a MAF of 0.129. For self-reported Hispanic Americans, the MAF of rs1555839 is 0.178.

* genome-wide significant (*p* < 5 × 10^−8^)

Due to significant correlations observed between RAN/RAS tasks and measures of reading fluency (Test of Word Reading Efficiency: TOWRE) and word reading (Woodcock-Johnson III Basic Reading; WJ-III Basic Reading), we wanted to evaluate whether allelic variation at rs1555839 (our most highly associated SNP in analyses of RAN/RAS) was associated with mean differences in performance for TOWRE and WJ-III Basic Reading (S1 Table). A univariate ANCOVA, covarying for the effects of age, sex, SES, and the first 10 principal components to correct for population stratification showed a significant effect of allele on TOWRE [F(1,1274)=13.15, p <0.001, η_p_^2^ = 0.01] and WJ-III Basic Reading [F(1,1271)=13.95, p <0.001, η_p_^2^ = 0.011] (Figure 2). Overall, performance on reading fluency and word reading was worse in the presence of the C allele at rs1555839, the same allele that was associated with reduced scores on RAN and RAS tests.

**Figure 2:**
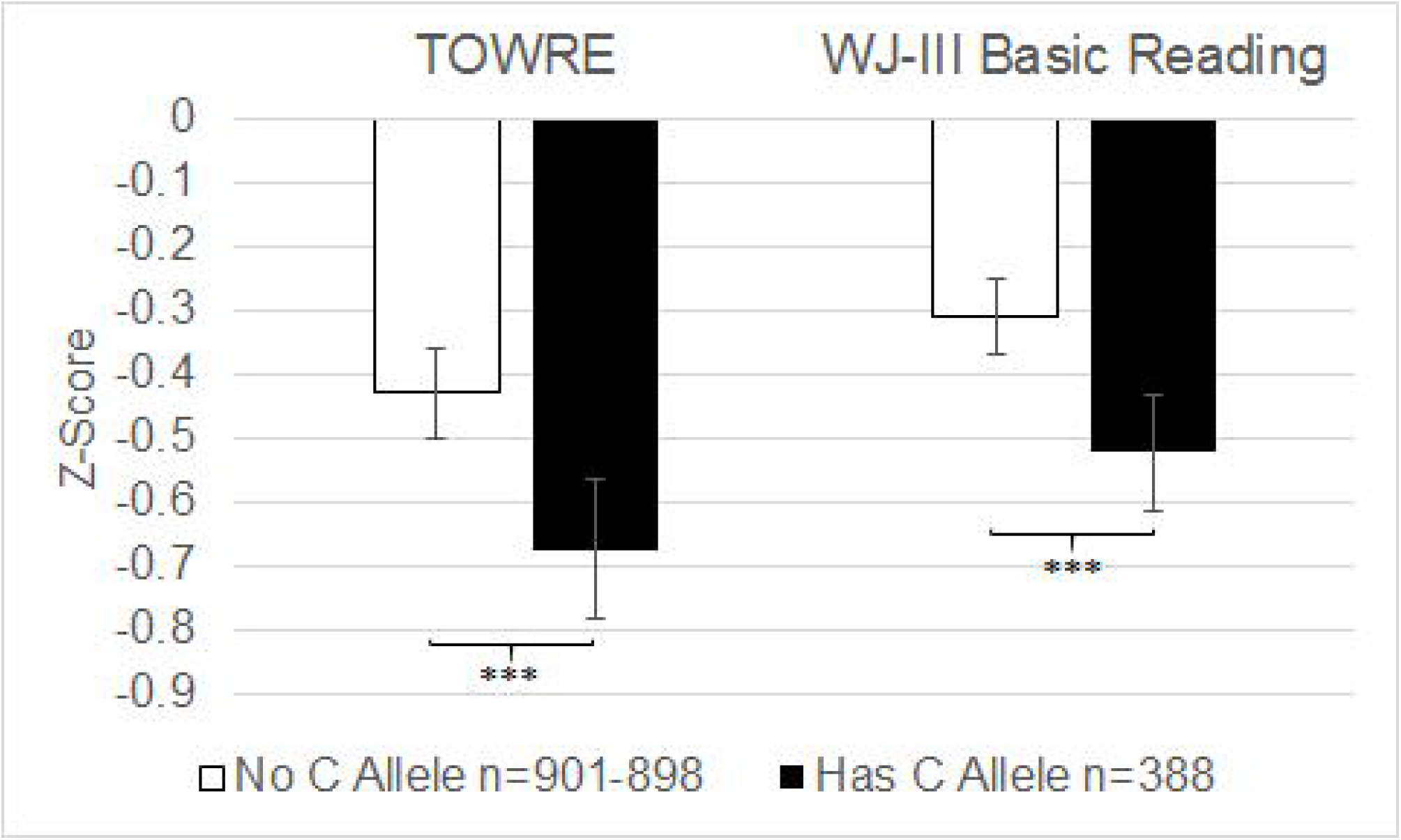
Mean differences in TOWRE and WJ-III Basic Reading performance by alleleic variation at rs1555839 in the GRaD Study. In the presence of the C allele there is a significant reduction in TOWRE and WJ-III Basic Reading performance, while correcting for the effects of age, sex, and population stratification (first 10 principal components). Error bars represent 95% confidence intervals. \*\*\**p* <0.001

In 318 unrelated subjects from the Colorado Learning Disabilities Research Center (CLDRC) cohort (ascertained based on RD and/or ADHD), RAN Colors, RAN Letters, RAN Numbers, and RAN Pictures were available for replication. RAN Objects and RAS Letters/Numbers were not collected in the CLDRC. Multivariate analysis showed replication of top markers within the pseudogene *RPL7P34* (Table 2). Specifically, rs10749593, rs7913742, rs1555839, and rs701825 were statistically significant (p < 0.05). Due to the non-independence of the most highly associated chromosome 10 markers in linkage disequilibrium (LD), no correction for multiple testing of SNPs was applied (S3 Figure). Post-hoc univariate association analysis of RAN Colors, RAN Letters, RAN Numbers, and RAN Pictures found a significant effect of rs7913742, rs1555839, and rs701825 across all RAN subtests, except for RAN Pictures (Table 2, S6 Table). In the CLDRC, consistent directions of effects on RAN Letters were observed with GRaD (S6 Table).

**Table 2:** CLDRC Multivariate Replication.

| MARKER | CHR | BP | Minor Allele | <sup>b</sup>MAF | p-value |  |  |  |  |
| --- | --- | --- | --- | --- | --- | --- | --- | --- | --- |
|  |  |  |  |  | Joint Model | RAN Pictures | RAN Colors | RAN Letters | RAN Numbers |
| rs2576167 | 10 | 90345671 | A | 0.302 | 0.067 | 0.592 | 0.051 | 0.027* | 0.064 |
| <sup>a</sup>rs1409136 | 10 | 90356598 | A | 0.303 | 0.061 | 0.568 | 0.052 | 0.025* | 0.064 |
| rs1359581 | 10 | 90360515 | A | 0.303 | 0.061 | 0.568 | 0.052 | 0.025* | 0.064 |
| rs10749593 | 10 | 90374057 | C | 0.319 | 0.025* | 0.416 | 0.052 | $5.44 \times 10^{-3}$* | 0.025* |
| <sup>a</sup>rs7913742 | 10 | 90376864 | C | 0.311 | 0.026* | 0.437 | 0.019* | $8.75 \times 10^{-3}$ | 0.024* |
| <sup>a</sup>rs1555839 | 10 | 90382820 | C | 0.319 | 0.019* | 0.410 | 0.034* | $5.21 \times 10^{-3}$* | 0.026* |
| <sup>a</sup>rs701825 | 10 | 90417547 | G | 0.282 | $2.47 \times 10^{-3}$* | 0.562 | $4.09 \times 10^{-3}$* | $9.56 \times 10^{-4}$* | $2.76 \times 10^{-3}$* |

<sup>a</sup> Imputed markers
\* $p < 0.05$ )

Results from the joint analysis of RAN Colors, RAN Pictures, RAN Letters, and RAN Numbers across the top associated markers in chromosome 10 of the GRaD discovery analysis. Results from post-hoc univariate analysis of RAN Pictures, RAN Colors, RAN Letters, and RAN Numbers at each marker are also represented. CHR = chromosome, BP = Base Position, MAF = Minor Allele Frequency

a Imputed markers

b MAF based on allele frequencies within the CLDRC sample

* p < 0.05)

### Bioinformatic Analysis

Using Genoskyline, we assessed whether the 70 kb region of chromosome 10q23.31 was a predicted functional region of the genome using high-throughput epigenomic annotations available through the NIH Epigenetic roadmap and ENCODE. Within the brain, Genoskyline predicted tissue specific functionality surrounding our top markers, with maximum GS scores of 1 (high probability that a region is associated with biological function) (Figure 3A). We then examined whether these findings could be attributed to epigenetic factors specific to a neural region. We found GS scores equal to 1 for predicted functionality in all tested regions of the brain surrounding *RNLS* (Figure 3A). Region specific functionality was observed in the cingulate gyrus, anterior caudate, hippocampus, and inferior temporal lobe within 2 kb of *RPL7P34* (Figure 3A).

**Figure 3:**
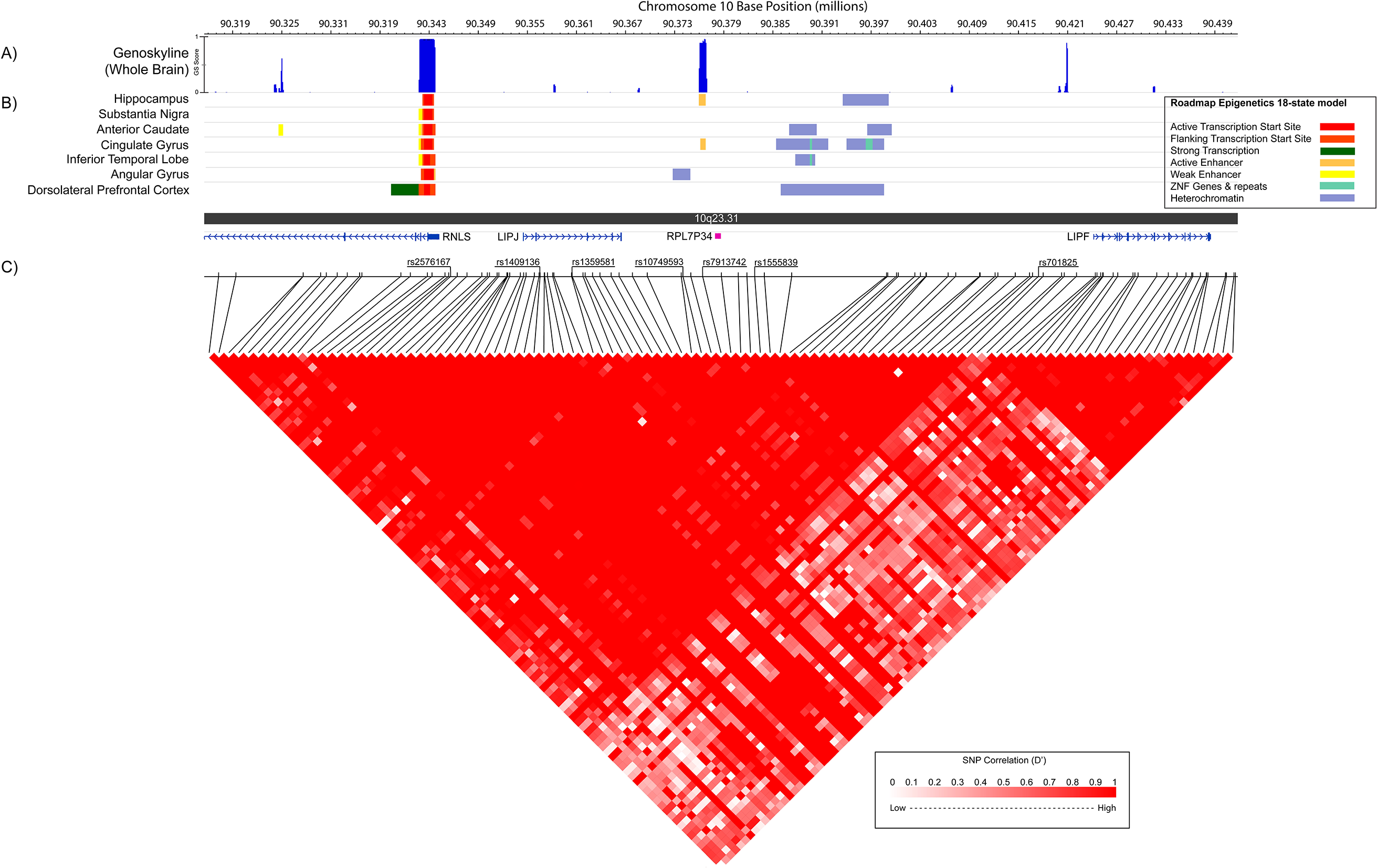
Bioinformatic examination of chromosome 10q23.31 containing the top performing markers in the GRaD discovery analysis. A) Plot of GS scores obtained from GenoSkyline indicating the posterior probability for functionality in the brain at each genomic locus. A GS score of 1 indicates a high probability for functionality, while a GS score of 0 suggests no functionality. B) 18-state chromatin model from Roadmap Epigenomics project showing predicted chromatin states based on the presence of H3K4me3, H3K4me1, H3K36me3, H3K27me3, H3K9me3, and H3K27ac sampled across different regions of the brain. C) Location of genes, markers assessed in the GRaD discovery analysis, and underlying LD structure (D’) across SNPs in the CEU population sampled in HapMap release 27.

To determine the chromatin state (i.e. weak enhancer region, active enhancer, active transcription start site, strong transcription) of the 70 kb region of chromosome 10q23.31 contributing to high posterior probability scores for tissue-specific functionality in the genome, we evaluated the 18-state model previously generated by the Roadmap Epigenomic Project. Briefly, a model was derived from a multivariate hidden Markov model that considers the combinatorial interactions of 6 histone marks (H3K4me3, H3K4me1, H3K36me3, H3K27me3, H3K9me3, and H3K27ac) across 127 epigenomes and predicts the chromatin state within a genomic region within 18 different classifications (35). Examination of the 18-state model revealed that the region closely associated with the most highly significant markers in the GRaD discovery analysis (rs1555839, rs10749593, rs7913742) is flanked by an active enhancer site in the hippocampus and cingulate gyrus, and regions of heterochromatin containing ZNF genes and repeats in the cingulate gyrus and inferior temporal lobe (Figure 3B). In addition, the 18-state model revealed an active transcription start site for *RNLS* in all evaluated brain regions (Figure 3B).

Examination of LD blocks across different ethnic groups sampled in the 1000 Genomes Project that are also represented in the GRaD sample (CEU, MEX, and YRI) show a single LD block spanning the 70 kb from rs2576167 to rs701825 (Figure 3C, S3 Figure), suggesting a similar underlying genomic architecture across ethnic groups (36).

### Pediatric, Imaging, Neurocognition, and Genetics (PING) Neuroimaging Genetic Analysis

Following replication of rs1555839, and bioinformatic analysis showing predicted functionality in brain tissue within the region of chromosome 10q23.31, we conducted a targeted neuroimaging genetics analysis in 690 typically developing children of various ethnic backgrounds from the PING sample, to determine whether rs1555839 was associated with variation in cortical volume of canonical left hemisphere regions of interest (ROIs) in the reading network and their right hemisphere counterparts. We found an association with right hemisphere inferior parietal cortex that survived correction for multiple testing(β = −432.3, p = 2.9 × 10^−3^; Table 3). The C allele was associated with lower cortical volumes in the right inferior parietal lobule in the PING sample.

**Table 3:** Reading-related regions of interest in the PING study

|  | LEFT |  |  | RIGHT |  |  |
| --- | --- | --- | --- | --- | --- | --- |
|  | BETA | SE | P | BETA | SE | P |
| Pars Opercularis | -19.33 | 67.07 | 0.7733 | 1.006 | 60.07 | 0.9866 |
| Pars Orbitalis | 14.54 | 26.82 | 0.588 | -15 | 32.15 | 0.6409 |
| Pars Triangularis | -62.3 | 52.4 | 0.2349 | -58.89 | 60.88 | 0.3337 |
| Supramarginal Gyrus | -166.8 | 120.1 | 0.1654 | 77.08 | 117.9 | 0.5133 |
| Inferior Parietal Cortex | -71.7 | 138.4 | 0.6046 | -432.3 | 144.4 | 0.0029* |
| Inferior Temporal Gyrus | -167.4 | 118.4 | 0.1576 | -30.86 | 119.7 | 0.7966 |
| Fusiform Gyrus | -100.6 | 95.21 | 0.2912 | -165.7 | 85.78 | 0.0538 |

Statistical models were corrected for general ancestry factor, age, sex, scanner device, handedness, intracranial volume, parental education, and family income.

* Survives Bonferroni correction for multiple testing (p < 0.05/14 = 0.0036)

## DISCUSSION

The present study is one of the first to examine the genetics of reading-related traits in an admixed population of Hispanic and African American children. Here, we described a multivariate GWAS that identified a region of chromosome 10q23.31 with pleiotropic effects across RAN Objects, RAN Letters, and RAS Letters/Numbers—all highly correlated tasks that are predictive of later reading outcome and reading ability in children and adults (14). Follow-up analysis in this same sample also showed that this region of chromosome 10 is associated with variation in tests for reading fluency (TOWRE) and word reading (WJ-III Basic Reading). Further bioinformatic analysis indicated that the most highly associated SNPs within chromosome 10q23.31 tag nearby regions of the genome with predicted function in the brain based on epigenetic markers. Additional neuroimaging genetics analysis in an independent sample revealed that rs1555839 is associated with structural variation in the right inferior parietal lobule. Intriguingly, variation in this region has been linked to RD in other studies (37, 38).

The most highly associated marker in the RAS/RAN genome-wide screen, rs1555839, is located upstream from the lncRNA pseudogene, *RPL7P34*. The function of *RPL7P34* is unknown, and the roles of lncRNAs in the genome are poorly understood. LncRNAs form a class of non-protein coding transcripts over 200 nucleotide bases long, and have characteristics that suggest functionality including tissue-specific expression, regulated expression, and regulation of gene expression and their networks (see (39) for review). LncRNAs have been reported to recruit transcription factors and interact with chromatin modifiers, suggesting that lncRNAs facilitate epigenetic regulation of the genome (40). Approximately 40% of lncRNAs are expressed in the brain and are hypothesized to play critical roles in neural development such as neural proliferation and differentiation (41).

In the present analysis, epigenetic examination of the region surrounding *RPL7P34* shows a predicted active enhancer site based on the presence of H3K4me1 and H3K27ac histone modifiers specifically in the brain. It is possible that *RPL7P34* may play a role in the recruitment of proteins that bind to enhancer sites, which promote transcription of nearby genes. The nearby gene *RNLS*, approximately 30 kb away, is the closest predicted transcription start site within the brain. The predicted enhancer and transcription start site lie within a topologically associating domain (TAD) identified in actively mitotic neural precursors in the developing brain (42). TADs refer to regions of the genome that physically interact more frequently, while physical interactions across two TADs are less likely to occur (43). In addition, these two sites are also in high linkage disequilibrium with each other across different ethnic populations relevant to this study (HapMap release 27: European, Mexican, and Yoruban; S3 Figure). Taken together, it is possible that the enhancer region downstream from *RPL7P34* could regulate *RNLS*. However, without further functional analyses, the molecular and epigenetic function of *RPL7P34* cannot be confirmed. It is also important to note that epigenetic data in the brain attained and analyzed by the Epigenetic Roadmap project were from only two individuals—one 75 and the other 81-years-old (35). Although these data offer some clues to potential functionality, there are limitations to their interpretation.

*RNLS* (also known as *C10orf59*) is a gene also located on chromosome 10q23.31 and implicated in our GWAS and bioinformatic analysis. *RNLS* encodes a flavin adenine dinucleotide-dependent amine oxidase, called renalase, which metabolizes catecholamines such as norepinephrine and dopamine—both of which are neurotransmitters produced by the brain, but also secreted by the adrenal glands to modulate autonomic responses (44, 45). Renalase is known to be secreted by the kidneys, circulates in blood, and modulates cardiac function and blood pressure (45). It is also present in the human central nervous system, specifically, the hypothalamus, pons, medulla oblongata, cerebellum, pituitary gland, cortex and spinal cord (44). *In vitro* analysis of monoamine oxidase activity of renalase shows that it is most efficient in metabolizing dopamine, but is also effective in metabolizing epinephrine and norepinephrine (45). In the brain, the dopaminergic and noradrenergic system are two major neuromodulatory systems that play important functions in motivation, attention, learning, and memory formation (46). Although bioinformatic evidence from the present study, expression data from GTEx and Brainspan, and evidence of renalase in *postmortem* human brain tissue support an active role in the brain, its functional role in metabolizing neurotransmitters in the brain is currently unclear (44). However, genetic variants in *RNLS* have been associated with schizophrenia, a neuropsychiatric disorder that may be caused by an imbalance in neurotransmission. Specifically, a recent study showed that human-induced pluripotent stem cell (hiPSC)-derived neurons from schizophrenia patients had altered catecholamine release relative to control hiPSC-derived neurons (47).

This is the first study to implicate *RNLS* in reading-related domains. However, it is not the first gene associated with the metabolism of catecholamines. *COMT* encodes catechol-O-methyltransferase, which degrades catecholamines in the brain, and has been associated with variation in reading-related tasks as well as functional networks associated with reading ability (48, 49). There is also evidence showing neurochemical differences between poor readers and typically developing controls suggesting that there could be alterations in how neurotransmitters are metabolized in reading impaired individuals (50).

Additional genes located in the region of chromosome 10q23.31 surrounding the top markers in the analysis include *LIPJ* and *LIPF*, which both encode proteins in the lipase family that are associated with lipid metabolism in the skin and stomach, respectively (51, 52). Both *LIPJ* and *LIPF* show no evidence for expression in the brain and it is unclear if they may have biological relevance in the etiology of RD.

Neuroimaging genetics analysis indicates that rs1555839 is associated with variation in cortical volume in the right inferior parietal cortex. The left inferior parietal cortex is part of the canonical reading network that includes the inferior frontal gyrus, temporoparietal region, and occipitotemporal area. However, in RD individuals, a more distributed network in brain activation across both left and right hemisphere structures during reading and RAN tasks are observed (14, 37, 38, 53). Limited studies have been conducted on structural neuroanatomical correlates of RAN, but there is evidence that RAN performance is associated with grey matter volumes in bilateral occipital-temporal and parietal-frontal regions, which include the right inferior parietal cortex (37). In addition, variation in the right inferior parietal cortex has also been implicated in number related tasks and may have implications for the rapid naming of numbers (54).

Previous genetic linkage studies for RAN have provided support for potential trait loci on chromosome 1p36-p34 (DYX8 locus) and 6p21 for a composite score of RAN Colors and Objects, 2p16-p15 (DYX3 locus) for RAN Letters, and 10q23.33-q24.32 for RAN Colors. The region most strongly implicated in the present analysis is on chromosome 10q23.31, which does not overlap with any of the above loci or other known genomic regions implicated in RD thus far. A potential reason why our results do not correspond with previously identified loci could be differences in the RAN data collected. In the GRaD study, the rapid naming tasks collected in the sample were RAN Objects, RAN Letters, and RAS Letters/Numbers. Grigorenko and colleagues (2001) and Konig and colleagues (2011) only showed linkage with RAN Colors, and neither included RAS Letters/Numbers. This suggests potential independent mechanisms associated with RAN Colors performance relative to RAN Objects, RAN Letters, and RAS Letters/Numbers (55). In addition, the above studies used linkage analyses, not association, to identify genetic regions linked to RAN and RAS performance—two distinct methods with different assumptions and sensitivities, which could explain the differences in results (56).

Most studies of the genetics of reading and RD have largely focused on populations of European descent. However, the genetic architecture of complex traits, like reading and related disorders may not share the same underlying genetics across ethnic groups. In fact, opposite effects of variants in the RD candidate risk gene *KIAA0319* have been reported between European and East Asian populations (57). Potential differences in the genetic architecture of RAN/RAS performance and RD across ethnic groups could also explain why the present results do not overlap with those previously identified. The study conducted by Rubenstein et al., (2015) is the closest in neurocognitive design to the present RAN/RAS analysis with the inclusion of a RAS subtest, but their sample largely consisted of individuals of European descent.

Small sample size is a limitation of this study for modern GWAS analyses. However, based on power calculations for a multivariate GWAS, we had moderate power of 0.68 to identify a genetic variant with a minor allele frequency of 0.16, effect size ranging from 0.03-0.013, and cross phenotype correlation of 0.7, in a sample with 1263 individuals (both phenotype and genotype data) and α = 5 × 10^−8^—all parameters applicable to this study and the observed SNPs showing most significant associations (58). While it is possible that the observed significant results could be attributed to population admixture, the calculated genomic inflation factor was within acceptable ranges (λ = 1.003-1.017) after correcting, suggesting adequate control for population stratification. An additional limitation of the study is that phenotypes used in the CLDRC replication sample did not perfectly match the RAN and RAS tasks evaluated in the GRaD sample. In the CLDRC cohort, performance on RAS Letters/Numbers was not collected. However, other rapid naming tasks, specifically RAN Colors and RAN Numbers, were available for analysis (RAN Letters and RAN Objects/Pictures overlapped). Although, RAS Letters/Numbers was not included in the CLDRC multivariate replication analysis, implicated SNPs in the chromosome 10 region were significant, providing support that the chromosome 10q23.31 had a pleiotropic effect across rapid naming tasks.

In conclusion, the current study identifies and replicates association of a region of chromosome 10q23.31 with pleiotropic effects across RAN, RAS, and reading abilities in a sample of Hispanic and African American youth. The present investigation leveraged different data sources and types across neuroimaging and epigenetic data. However, further functional assays must be conducted to shed mechanistic light on the pathways involved. There is growing evidence that noncoding regions of the genome have an impact on reading-related traits, and the identification of a lncRNA associated with a reading endophenotype lends additional support (59). Implication of *RNLS* also reinforces the hypothesis that alterations in neurochemical modulation in the brain could contribute to impairments in reading performance. Lastly, this study highlights the importance of studying the genetic architecture of RD across diverse ethnicities and how genetic effects and variants may differ (or be similar) across populations. This is not only critical in our understanding of the biological mechanisms that contribute to RD, but also necessary for presymptomatic identification and development of precision intervention strategies informed by genetic screening.

## MATERIALS AND METHODS

### Genes, Reading, and Dyslexia Study

The Genes, Reading, and Dyslexia (GRaD) study, is a multisite case-control study of RD in minority youth across the United States, Canada, and Puerto Rico. Detailed descriptions of recruitment, inclusion, and exclusion criteria for the sample are reported elsewhere (60). Briefly, male and female children age 8-15 years of African American and/or Hispanic American ancestry were recruited for study (S2 Table). Children were excluded if they were not of African American or Hispanic American ancestry, placed in foster care, had a medical history or neurological condition that could affect cognitive or neural development (i.e. preterm birth, prolonged stay in the NICU, seizures, acquired brain injuries), diagnosis of any cognitive or neuropsychiatric disorder (i.e. intellectual disability, autism spectrum disorder, depression), or documented hearing or vision impairment. Inclusion criteria for participants likely to have a reading disability included either history of poor reading skills (as documented by prior school or clinical testing), report of skills falling below expected level for age or grade, and/or provision of special services in the area of reading. For inclusion of participants likely to be controls, inclusion criteria were “competent reading skills” as identified by reading skills falling at or above current expectations for grade and performance falling above the 40th percentile on standardized school or clinical testing. A total of 1,432 unrelated children were recruited into the GRaD study. Of these subjects, 1,331 children with high-quality DNA samples were included in the analysis. Informed consent was obtained for all participants and parents or legal guardians of subjects. Ethical approval of study protocols and recruitment was obtained by Institutional Review Boards at each recruiting site (University of Colorado-Boulder, University of Denver, Tufts University, University of New Mexico, Kennedy Krieger Institute, Hospital for Sick Children-Toronto, and Yale University).

### Rapid Automatized Naming (RAN) and Rapid Automatized Stimulus Measures (RAS)

RAN and RAS performance in the GRaD sample was evaluated using the RAN Objects, RAN Letters, and RAS Letters/Numbers tasks developed by Wolf and Denckla (61). For RAN objects and RAN letters, subjects name aloud 50 familiar, high frequency objects or letters, respectively, arranged in a 5 × 10 array as quickly and accurately as possible. The format of RAS Letters/Numbers is similar to RAN Objects and RAN Letters, except that items in the array are from alternating stimulus categories (e.g. letters and numbers). Time to complete each task is recorded and converted to age-standardized scores. Age-standardized scores (mean of 100, SD of 15) were then converted to Z-scores (mean of 0, SD of 1) used for downstream genetic analyses (S7 Table).

### Test of Word Reading Efficiency (TOWRE)

The TOWRE Total Word Reading Efficiency is a composite of both Sight Word Efficiency and Phonetic Decoding Efficiency scores and is an assessment of reading fluency (the ability to read words quickly and accurately) under timed conditions (62). For this assessment, the subject is evaluated on the number of individual words (Sight Word Efficiency) and nonwords (Phonetic Decoding Efficiency) correctly read in 45 seconds. For each subtest, the total number of words read correctly is converted into a standard score based on age norms and then z-scored (S7 Table). The TOWRE Total Word Reading Efficiency composite was used for downstream analysis.

### Woodcock–Johnson Tests of Achievement, Third Edition (WJ-III)

The WJ-III Basic Reading Score is a composite of the WJ-III Letter-Word Identification and WJ-III Word Attack subtests (63). The WJ-III Letter Word Identification subtest is an untimed measure of reading increasingly complex English words aloud. The Word Attack subtest is a decoding measure of nonwords or pseudowords in isolation. For each subtest, the total number of words read correctly is converted into a standard score based on age norms and then z-scored (S7 Table). The WJ-III Basic Reading composite was used for downstream analysis.

### DNA Collection, Genotyping, and Analysis

Saliva was collected from each GRaD subject using the Oragene-DNA self-collection kit (OG-500; DNA Genotek Inc, Ottawa, Ontario, Canada). DNA was extracted using prepIT-L2P (DNA Genotek Inc, Ottawa, Ontario, Canada). Subjects were successfully genotyped for 2,391,739 single nucleotide polymorphisms (SNPs) using the Illumina Infinium Omni2.5-8 BeadChip at the Yale Center for Genome Analysis (Orange, CT). Initial genotyping quality control and SNP genotype calls were conducted using GenomeStudio (Illumina, San Diego, CA) and standard Infinium genotyping data analysis parameters to optimize genotyping accuracy. SNPs were removed from downstream analysis if they had missingness greater than 5% (n=22,849), Hardy-Weinberg equilibrium p<0.0001 (n=116,259), were not autosomal (n=60,551), or had a minor allele frequency less than 10% (n=1,182,060). Samples were removed if they were missing more than 3% of their genotypes (n=39), if there were discrepancies between reported and inferred sex based on X chromosome heterozygosity (n=52), and IBD > 0.125 calculated using REAP (n=10) (64). After quality control, there were a total 1,331 samples genotyped with 1,010,020 SNPs.

#### Population Stratification

The first 10 principal components derived from genome-wide SNP data were used to correct for genomic inflation due to allele frequency differences across different ancestries (population stratification) with EIGENSTRAT (65). Population stratification was evaluated using a genomic inflation factor (λ) calculated using PLINK (66). A λ factor below the standard threshold of 1.05 indicates sufficient correction for population stratification. Since calculated λ ranged from 1.003-1.017 in the present study, 10 principal components were sufficient to correct for population stratification (S1 Figure).

#### Statistical Analysis

Univariate genetic analyses for RAN objects, RAN letters, and RAS Letters/Numbers were performed using PLINK v1.9 to test each SNP using linear regression under an additive model. Multivariate genetic analysis jointly analyzing RAN objects, RAN letters, and RAS Letters/Numbers for pleiotropic effects was conducted using the R package MultiPhen (67). MultiPhen allows for the simultaneous examination of multiple correlated traits using a reversed ordinal regression to determine the linear combination of traits most associated with specific genotypes at each SNP. It then performs a log likelihood ratio test on the joint model against the null model to evaluate association. All statistical models were corrected for the first 10 principal components to correct for population stratification, sex, age, and socioeconomic status (SES). In this study, SES was assessed as a binary variable that describes whether the subject is enrolled in at least 1 government assistance program with an income eligibility requirement (e.g. food stamps, Medicaid, housing choice voucher program, and/or Women, Infants, and Children program). To correct for multiple testing, we used the standard threshold of 5 × 10^−8^ (Bonferroni correction for 1 million tests) to determine genome-wide significance.

#### Replication Analysis

### Colorado Learning Disability Research Center (CLDRC) Cohort

Replication was conducted on samples from the CLDRC. Methods related to recruitment, ascertainment, data collection (neurocognitive and genetic), and data processing are described in detail elsewhere (10, 68). Briefly, the CLDRC sample is a selected twin cohort for RD, ADHD, and other learning disabilities recruited from 27 school districts in Colorado (68, 69). Subjects were assessed with RAN colors, RAN objects, RAN letters, and RAN numbers (70). For this task, participants named as many items in a 15 x 5 array as quickly and accurately as possible. The number of correctly named items in 15 seconds was recorded. Raw scores were standardized and age-regressed for each of the tasks based on a separate control sample consisting of typically developing children. All experimental procedures and written informed consent forms were approved by the institutional review boards (IRB) at University of Colorado-Boulder and University of Denver.

DNA was collected and extracted from saliva and genotyped using the Illumina Human OmniExpress genotyping panel (713,599 SNPs). Initial genotyping quality control and SNP genotype calls were conducted using GenomeStudio (Illumina, San Diego, CA). Initial quality control filters included the removal of samples with a call rate <98% and SNPs with a call rate <95%, HWE <0.0001, and MAF < 5%. SNPs on chromosome 10 identified in the GRaD discovery analysis that were not genotyped were imputed with genipe—an automated genome-wide imputation pipeline that executes PLINK 1.07 (66), SHAPEIT (71), and IMPUTE2 (72) for data imputation to the 1000 Genomes Project, Phase 3 reference (36, 73). All imputed SNPs had an info score (IMPUTE2 imputation quality metric) greater than 0.9, indicating that all replication SNPs were imputed with high confidence.

The sample of twins and siblings available for this study comprised 749 participants in total, mean age 11.7 years, age range 8–19, from 343 unrelated twinships/sibships. For the present study, only participants of European descent were analyzed and one child per twinship/sibship were randomly selected for analysis based on the availability of RAN and genetic data. The total sample size for replication analysis was 318 unrelated individuals.

Multivariate genetic analysis jointly analyzing RAN colors, RAN objects, RAN letters, and RAN numbers standard scores for association at candidate chromosome 10 SNPs identified in the GRaD discovery analysis was conducted using MultiPhen while covarying for the effects of age and sex.

#### Bioinformatic Analysis

GenoSkyline is an unsupervised learning framework that predicts tissue-specific functionality in non-coding regions of the genome by integrating genome-wide epigenetic data from the Roadmap Epigenomics Project and ENCODE (35, 74, 75). Detailed description of tissue specific functionality has been previously described (74). Briefly, GenoSkyline uses an unsupervised-learning technique that evaluates the presence of well characterized histone marks (H3k4me1, H3k4me3, H3k36me3, H3k27me3, H3k9me3, H3k27ac, H3k9ac) and DNase 1 hypersensitivity sites and calculates a posterior probability score (GS score) that a given genetic coordinate is functional. A GS score of “1” suggests that the genomic region of interest is functional within the given tissue type, while a score of “0” suggests no functional significance. Pre-calculated, genome-wide, tissue-specific GS scores for blood, epithelium, muscle, heart, lung, gastrointestinal, brain, and sub-regions of the brain (angular gyrus, prefrontal cortex, cingulate gyrus, anterior caudate, hippocampus, inferior temporal gyrus, and substantia nigra) were obtained from the GenoSkyline database (<u>http://genocanyon.med.yale.edu/GenoSkyline</u>). Follow-up analysis of epigenetic data from the Roadmap epigenetics project was conducted to identify predicted chromatin state in the genomic region of interest (<u>http://egg2.wustl.edu/roadmap/web_portal/index.html</u>). Data were visualized using the Washington University in St. Louis (WashU) EpiGenome Browser v.42.

#### Neuroimaging Genetic Analysis

### Pediatric, Imaging, Neurocognition, and Genetics (PING) Study

Neuroimaging genetics analysis of volumes of cortical ROIs was conducted in the PING sample (http://ping.chd.ucsd.edu/). Methods related to recruitment, ascertainment, data collection (neuroimaging, neurocognitive, genetic), and data processing are described in detail elsewhere (76). Briefly, the PING study is a cross sectional sample of typically developing children ranging in age from 3-20 years old. Individuals were excluded from participating if they had a history of major developmental, psychiatric, or neurological disorders, brain injury, prematurity (i.e., born at less than 36 weeks gestational age), prenatal exposure to illicit drugs or alcohol, history of head trauma, or other medical conditions that could affect development. Subjects with learning disability or ADHD were not excluded. All experimental procedures and written informed consent forms were approved by the institutional review boards (IRB) at each of the 10 participating PING study recruitment sites (University of California at San Diego, University of Hawaii, University of California at Los Angeles, Children's Hospital of Los Angeles of the University of Southern California, University of California at Davis, Kennedy Krieger Institute of Johns Hopkins University, Sackler Institute of Weill Cornell Medical College, University of Massachusetts, Massachusetts General Hospital at Harvard University, and Yale University). Parental informed consent was obtained for participants less than 18 years of age with child assent (ages 7-17). For individuals 18 years of age and older, written informed consent was obtained.

DNA was extracted from saliva and genotyped on the Illumina Human660W Quad BeadChip (655,214 SNPs) for 1391 subjects in the PING study. Initial genotyping quality control and SNP genotype calls was conducted using GenomeStudio (Illumina, San Diego, CA) by the PING genomics core at the Scripps Translational Science Institute (La Jolla, CA). Initial quality control filters included the removal of samples with a call rate <98% and SNPs with a call rate <95%, HWE <0.0001, and MAF < 5%. To assess ancestry and admixture proportions in the PING participants, a supervised clustering approach implemented in the ADMIXTURE software grouped participants into six clusters corresponding to six major continental populations: African, Central Asian, East Asian, European, Native American and Oceanic (77). Implementation of ancestry and admixture proportions in the PING subjects is described in detail elsewhere (76). To prevent possible population stratification, genetic ancestry factor (GAF) was included as a covariate in all analyses.

In depth descriptions of methods for neuroimaging data acquisition and processing for the PING study are described elsewhere (76). Briefly, structural MRI data were collected from all individuals using a standardized multiple-modality high-resolution structural MRI protocol involving 3D T1-weighted volumes across sites to maintain consistency across data collection. Image files in DICOM format were processed by the neuroimaging post-processing core at the University of California at San Diego using an automated processing stream written in MATLAB and C++. Cortical surface reconstruction and subcortical segmentation were performed using a fully automated set of tools available in the Freesurfer software suite (<u>http://surfer.nmr.mgh.harvard.edu/</u>). Cortical parcellation of sulci and gyri were automatically defined using the Desikan-Killiany Atlas integrated within the FreeSurfer suite (78). All data (genetic, neuroanatomical, and cognitive) were obtained from the PING portal (<u>https://ping-xdataportal.ucsd.edu/</u>).

### Cortical volume regions of interest selection and statistical analysis

Left and right hemisphere cortical regions spanning the inferior frontal gyrus, temporoparietal region, and occipitotemporal area were selected for candidate region of interest analysis (S8 Table). Left hemisphere structures spanning the inferior frontal gyrus, temporoparietal region, and occipitotemporal area comprise the canonical reading network, and show atypical patterns of functional activation in reading disabled children relative to typically developing children (38). Respective right hemisphere regions were also included in the analysis since evidence suggests that naming speed is also associated with grey matter volume differences in right hemisphere frontal, temporoparietal and occipital regions (14, 38).

Imaging genetics analysis was conducted on 690 subjects with complete phenotype and genotype data that passed both neuroimaging and genotype QC. All analyses were corrected for the effects of age, sex, handedness, scanner device (79), genetic ancestry (African, Central Asian, East Asian, European, Native American and Oceanic), intracranial volume, highest parental education, and family income. The marker rs1555839 was genotyped using the Illumina Human660W Quad BeadChip in the PING sample and used for neuroimaging genetics analysis. Cortical volumes across 14 ROIs were tested for association with rs1555839 using linear regression under an additive genetic model in PLINK 1.9 (80).

## ACKNOWLEDGEMENTS

We extend our sincere gratitude to all the individuals and their families who participated in this study, and all the research assistants for their help with recruitment and data collection. We thank the staff at the Yale Center for Genome Analysis genotyping services. We also thank Dr. Mellissa DeMille, Dr. Jeffrey Malins, and Dr. Chintan Mehta for invaluable discussion and critical evaluation of this manuscript.

## FUNDING

The Genes, Reading, and Dyslexia Study was funded by The Manton Foundation (JRG). The Colorado Learning Disabilities Cohort was funded by the Eunice Kennedy Shriver National Institute of Child Health & Human Development National Institutes of Health Grant P50HD027802 (JRG, JCD, RKO, BFP, SSD, EGW). The Pediatric, Imaging, Neurocognition, and Genetics Study was funded by the National Institute on Drug Abuse and the Eunice Kennedy Shriver National Institute of Child Health & Human Development grant RC2DA029475. DTT and AKK were each funded by a training grant from the Eunice Kennedy Shriver National Institute of Child Health & Human Development 5T32HD07094 and 5T32HD007149, respectively. The Max Planck Society supported AG, CF and SEF, and funded the genetic analyses of the CLDRC cohort. The funders had no role in study design, data collection and analysis, decision to publish, or preparation of the manuscript.

## SUPPLEMENTAL TABLES LEGENDS

**S1 Table: Pearson correlation coefficients across RAN Objects, RAN Letters, RAS Letters/Numbers, Test of Word Reading Efficiency (TOWRE), and Woodcock-Johnson Basic Reading (WJ-III Basic Reading).** **p<0.01

**S2 Table: GRaD Demographics.** Demographic information on 1,331 children with available genetic data in the GRaD sample. ^a^ Enrolled in one or more government assistance programs including food stamps, Medicaid, housing choice voucher program, and women, infants, and children program.

**S3 Table: Post-hoc univariate examination of RAS Letters/Numbers across the top associated markers in the multivariate RAN/RAS analysis in GRaD.**

**S4 Table: Post-hoc univariate examination of RAN Objects across the top associated markers in the multivariate RAN/RAS analysis in GRaD.**

**S5 Table: Post-hoc univariate examination of RAN Letters across the top associated markers in the multivariate RAN/RAS analysis in GRaD.**

**S6 Table: Univariate association analysis of RAN Letters in the CLDRC cohort displaying results from top chromosome 10 markers identified in the GRaD Multivariate discovery GWAS.** *p<0.05

**S7 Table: GRaD Assessments Descriptives:** Descriptive statistics for assessments evaluated in the GRaD sample. Means reflect the z-score (mean = 0, standard deviation (SD) = 1) of the population standard score (mean = 100, SD = 15) available for each psychometric analysis

**S8 Table: Cortical regions of interest associated with reading ability and disability**

## SUPPLEMENTAL FIGURES LEGENDS

**S1 Figure: Plot of the first three principal components generated from genome-wide SNP data used to correct for population stratification.** The plot displays the clustering of individuals across the first three principal components and its correspondence to self-reported ancestry. Colors represent self-reported ancestry.

**S2 Figure: Manhattan plot representing the −log_10_*p* at each SNP assessed in the Multivariate analysis of RAN Objects, RAN Letters, and RAN Letters/Numbers.** Red line indicates the standard threshold for genome-wide significance (*p* < 5 × 10^−8^).

**S3 Figure: LD heatmaps across CEU, MXL, and YRI populations in the region of chromosome 10q23.31 containing the top performing markers in the GRaD discovery analysis.**

## REFERENCES

1. Shaywitz SE, Shaywitz BA. Dyslexia (Specific Reading Disability). Biological psychiatry. 2005;57(11):1301–9.

2. Hensler BS, Schatschneider C, Taylor J, Wagner RK. Behavioral genetic approach to the study of dyslexia. Journal of Developmental and Behavioral Pediatrics. 2010;31(7):525–32.

3. Logan JAR, Hart SA, Cutting L, Deater-Deckard K, Schatschneider C, Petrill S. Reading Development in Young Children: Genetic and Environmental Influences. Child Development. 2013;84(6):2131–44.

4. Carrion-Castillo A, Franke B, Fisher SE. Molecular Genetics of Dyslexia: An Overview. Dyslexia. 2013;19(4):214–40.

5. Meng H, Smith SD, Hager K, Held M, Liu J, Olson RK, et al. DCDC2 is associated with reading disability and modulates neuronal development in the brain. Proceedings of the National Academy of Sciences of the United States of America. 2005;102(47):17053–8.

6. Powers Natalie R, Eicher John D, Butter F, Kong Y, Miller Laura L, Ring Susan M, et al. Alleles of a Polymorphic ETV6 Binding Site in DCDC2 Confer Risk of Reading and Language Impairment. The American Journal of Human Genetics. 2013;93(1):19–28.

7. Plomin R. Child development and molecular genetics: 14 years later. Child Dev. 2013;84(1):104–20.

8. Luciano M, Evans DM, Hansell NK, Medland SE, Montgomery GW, Martin NG, et al. A genome-wide association study for reading and language abilities in two population cohorts. Genes Brain Behav. 2013;12(6):645–52.

9. Eicher JD, Powers NR, Miller LL, Akshoomoff N, Amaral DG, Bloss CS, et al. Genome-wide association study of shared components of reading disability and language impairment. Genes, Brain and Behavior. 2013;12(8):792–801.

10. Gialluisi A, Newbury DF, Wilcutt EG, Olson RK, DeFries JC, Brandler WM, et al. Genome-wide screening for DNA variants associated with reading and language traits. Genes Brain Behav. 2014;13(7):686–701.

11. Glahn DC, Knowles EE, McKay DR, Sprooten E, Raventos H, Blangero J, et al. Arguments for the sake of endophenotypes: examining common misconceptions about the use of endophenotypes in psychiatric genetics. Am J Med Genet B Neuropsychiatr Genet. 2014;165B(2):122–30.

12. Gershon ES, Goldin LR. Clinical methods in psychiatric genetics. I. Robustness of genetic marker investigative strategies. Acta Psychiatr Scand. 1986;74(2):113–8.

13. Gottesman, II, Gould TD. The endophenotype concept in psychiatry: etymology and strategic intentions. Am J Psychiatry. 2003;160(4):636–45.

14. Norton ES, Wolf M. Rapid automatized naming (RAN) and reading fluency: implications for understanding and treatment of reading disabilities. Annu Rev Psychol. 2012;63:427–52.

15. Denckla MB, Rudel R. Rapid “automatized” naming of pictured objects, colors, letters and numbers by normal children. Cortex. 1974;10(2):186–202.

16. Waber DP, Forbes PW, Wolff PH, Weiler MD. Neurodevelopmental characteristics of children with learning impairments classified according to the double-deficit hypothesis. J Learn Disabil. 2004;37(5):451–61.

17. Wolf M, O’Rourke AG, Gidney C, Lovett M, Cirino P, Morris R. The second deficit: An investigation of the independence of phonological and naming-speed deficits in developmental dyslexia. Read Writ. 2002;15(1):43–72.

18. Katzir T, Kim YS, Wolf M, Morris R, Lovett MW. The varieties of pathways to dysfluent reading: comparing subtypes of children with dyslexia at letter, word, and connected text levels of reading. J Learn Disabil. 2008;41(1):47–66.

19. Torppa M, Lyytinen P, Erskine J, Eklund K, Lyytinen H. Language Development, Literacy Skills, and Predictive Connections to Reading in Finnish Children With and Without Familial Risk for Dyslexia. Journal of Learning Disabilities. 2010;43(4):308–21.

20. Schatschneider C, Fletcher JM, Francis DJ, Carlson CD, Foorman BR. Kindergarten Prediction of Reading Skills: A Longitudinal Comparative Analysis. Journal of Educational Psychology. 2004;96(2):265–82.

21. Vukovic RK, Wilson AM, Nash KK. Naming speed deficits in adults with reading disabilities: a test of the double-deficit hypothesis. J Learn Disabil. 2004;37(5):440–50.

22. van den bos kp, Zijlstra BJH, Spelberg HC. Life-Span Data on Continuous-Naming Speeds of Numbers, Letters, Colors, and Pictured Objects, and Word-Reading Speed. Scientific Studies of Reading 2002;6(1):25–49.

23. Wolf M. Rapid alternating stimulus naming in the developmental dyslexias. Brain Lang. 1986;27(2):360–79.

24. Altemeier LE, Abbott RD, Berninger VW. Executive functions for reading and writing in typical literacy development and dyslexia. J Clin Exp Neuropsychol. 2008;30(5):588–606.

25. Davis CJ, Gayan J, Knopik VS, Smith SD, Cardon LR, Pennington BF, et al. Etiology of reading difficulties and rapid naming: the Colorado Twin Study of Reading Disability. Behav Genet. 2001;31(6):625–35.

26. Petrill SA, Thompson LA, Deater-Deckard K, Dethorne LS, Schatschneider C. Genetic and Environmental Effects of Serial Naming and Phonological Awareness on Early Reading Outcomes. J Educ Psychol. 2006;98(1):112–21.

27. Grigorenko EL, Wood FB, Meyer MS, Pauls JE, Hart LA, Pauls DL. Linkage studies suggest a possible locus for developmental dyslexia on chromosome 1p. Am J Med Genet. 2001;105(1):120–9.

28. Konig IR, Schumacher J, Hoffmann P, Kleensang A, Ludwig KU, Grimm T, et al. Mapping for dyslexia and related cognitive trait loci provides strong evidence for further risk genes on chromosome 6p21. Am J Med Genet B Neuropsychiatr Genet. 2011;156B(1):36–43.

29. Rubenstein KB, Raskind WH, Berninger VW, Matsushita MM, Wijsman EM. Genome scan for cognitive trait loci of dyslexia: Rapid naming and rapid switching of letters, numbers, and colors. Am J Med Genet B Neuropsychiatr Genet. 2014;165B(4):345–56.

30. de Kovel CG, Franke B, Hol FA, Lebrec JJ, Maassen B, Brunner H, et al. Confirmation of dyslexia susceptibility loci on chromosomes 1p and 2p, but not 6p in a Dutch sib-pair collection. Am J Med Genet B Neuropsychiatr Genet. 2008;147(3):294–300.

31. Carrion-Castillo A, Maassen B, Franke B, Heister A, Naber M, van der Leij A, et al. Association analysis of dyslexia candidate genes in a Dutch longitudinal sample. Eur J Hum Genet. 2017;25(4):452–60.

32. Lim CK, Wong AM, Ho CS, Waye MM. A common haplotype of KIAA0319 contributes to the phonological awareness skill in Chinese children. Behav Brain Funct. 2014;10:23.

33. Carrion-Castillo A, van Bergen E, Vino A, van Zuijen T, de Jong PF, Francks C, et al. Evaluation of results from genome-wide studies of language and reading in a novel independent dataset. Genes Brain Behav. 2016;15(6):531–41.

34. Galesloot TE, van Steen K, Kiemeney LALM, Janss LL, Vermeulen SH. A Comparison of Multivariate Genome-Wide Association Methods. PLOS ONE. 2014;9(4):e95923.

35. Roadmap Epigenomics C, Kundaje A, Meuleman W, Ernst J, Bilenky M, Yen A, et al. Integrative analysis of 111 reference human epigenomes. Nature. 2015;518(7539):317–30.

36. Genomes Project C, Auton A, Brooks LD, Durbin RM, Garrison EP, Kang HM, et al. A global reference for human genetic variation. Nature. 2015;526(7571):68–74.

37. He Q, Xue G, Chen C, Chen C, Lu ZL, Dong Q. Decoding the neuroanatomical basis of reading ability: a multivoxel morphometric study. J Neurosci. 2013;33(31):12835–43.

38. Norton ES, Beach SD, Gabrieli JD. Neurobiology of dyslexia. Curr Opin Neurobiol. 2015;30:73–8.

39. Wu H, Yang L, Chen L-L. The Diversity of Long Noncoding RNAs and Their Generation. Trends in Genetics.

40. Smith JE, Alvarez-Dominguez JR, Kline N, Huynh NJ, Geisler S, Hu W, et al. Translation of small open reading frames within unannotated RNA transcripts in Saccharomyces cerevisiae. Cell Rep. 2014;7(6):1858–66.

41. Roberts TC, Morris KV, Wood MJ. The role of long non-coding RNAs in neurodevelopment, brain function and neurological disease. Philos Trans R Soc Lond B Biol Sci. 2014;369(1652).

42. Won H, de la Torre-Ubieta L, Stein JL, Parikshak NN, Huang J, Opland CK, et al. Chromosome conformation elucidates regulatory relationships in developing human brain. Nature. 2016;538(7626):523–7.

43. Dixon JR, Selvaraj S, Yue F, Kim A, Li Y, Shen Y, et al. Topological domains in mammalian genomes identified by analysis of chromatin interactions. Nature. 2012;485(7398):376–80.

44. Hennebry SC, Eikelis N, Socratous F, Desir G, Lambert G, Schlaich M. Renalase, a novel soluble FAD-dependent protein, is synthesized in the brain and peripheral nerves. Mol Psychiatry. 2010;15(3):234–6.

45. Xu J, Li G, Wang P, Velazquez H, Yao X, Li Y, et al. Renalase is a novel, soluble monoamine oxidase that regulates cardiac function and blood pressure. J Clin Invest. 2005;115(5):1275–80.

46. Kobayashi K. Role of catecholamine signaling in brain and nervous system functions: new insights from mouse molecular genetic study. J Investig Dermatol Symp Proc. 2001;6(1):115–21.

47. Hook V, Brennand KJ, Kim Y, Toneff T, Funkelstein L, Lee KC, et al. Human iPSC neurons display activity-dependent neurotransmitter secretion: aberrant catecholamine levels in schizophrenia neurons. Stem Cell Reports. 2014;3(4):531–8.

48. Landi N, Frost SJ, Mencl WE, Preston JL, Jacobsen LK, Lee M, et al. The COMT Val/Met polymorphism is associated with reading-related skills and consistent patterns of functional neural activation. Dev Sci. 2013;16(1):13–23.

49. Grigorenko EL, Deyoung CG, Getchell M, Haeffel GJ, Klinteberg BA, Koposov RA, et al. Exploring interactive effects of genes and environments in etiology of individual differences in reading comprehension. Dev Psychopathol. 2007;19(4):1089–103.

50. Pugh KR, Frost SJ, Rothman DL, Hoeft F, Del Tufo SN, Mason GF, et al. Glutamate and choline levels predict individual differences in reading ability in emergent readers. J Neurosci. 2014;34(11):4082–9.

51. Lohse P, Lohse P, Chahrokh-Zadeh S, Seidel D. The acid lipase gene family: three enzymes, one highly conserved gene structure. J Lipid Res. 1997;38(5):880–91.

52. Toulza E, Mattiuzzo NR, Galliano MF, Jonca N, Dossat C, Jacob D, et al. Large-scale identification of human genes implicated in epidermal barrier function. Genome Biol. 2007;8(6):R107.

53. Misra M, Katzir T, Wolf M, Poldrack RA. Neural Systems for Rapid Automatized Naming in Skilled Readers: Unraveling the RAN-Reading Relationship. Scientific Studies of Reading. 2004;8(3):241–56.

54. Hoeft F, Meyler A, Hernandez A, Juel C, Taylor-Hill H, Martindale JL, et al. Functional and morphometric brain dissociation between dyslexia and reading ability. Proc Natl Acad Sci U S A. 2007;104(10):4234–9.

55. Tannock R, Martinussen R, Frijters J. Naming speed performance and stimulant effects indicate effortful, semantic processing deficits in attention-deficit/hyperactivity disorder. J Abnorm Child Psychol. 2000;28(3):237–52.

56. Borecki IB. Linkage and Association Studies. eLS: John Wiley & Sons, Ltd; 2001.

57. Shao S, Niu Y, Zhang X, Kong R, Wang J, Liu L, et al. Opposite Associations between Individual KIAA0319 Polymorphisms and Developmental Dyslexia Risk across Populations: A Stratified Meta-Analysis by the Study Population. Sci Rep. 2016;6:30454.

58. Porter HF, O’Reilly PF. Multivariate simulation framework reveals performance of multitrait GWAS methods. Sci Rep. 2017;7:38837.

59. Devanna P, Chen XS, Ho J, Gajewski D, Smith SD, Gialluisi A, et al. Next-gen sequencing identifies non-coding variation disrupting miRNA-binding sites in neurological disorders. Mol Psychiatry. 2017.

60. Jacobson LA, Koriakin T, Lipkin P, Boada R, Frijters JC, Lovett MW, et al. Executive Functions Contribute Uniquely to Reading Competence in Minority Youth. J Learn Disabil. 2016.

61. Wolf M, Denkla MB. RAN/RAS: Rapid Automatized Naming and Rapid Alternating Stimulus Tests. Austin, TX: Pro-Ed; 2005.

62. Torgesen JK, Wagner RK, Rashotte CA. Test of Word Reading Efficiency. Austin, TX: PRO-ED; 1999.

63. Woodcock RW, McGrew KS, Mather N. Woodcock-Johnson^®^ III Normative Update Complete. Boston, MA: Houghton Mifflin Harcourt; 2001.

64. Thornton T, Tang H, Hoffmann TJ, Ochs-Balcom HM, Caan BJ, Risch N. Estimating kinship in admixed populations. Am J Hum Genet. 2012;91(1):122–38.

65. Price AL, Patterson NJ, Plenge RM, Weinblatt ME, Shadick NA, Reich D. Principal components analysis corrects for stratification in genome-wide association studies. Nat Genet. 2006;38(8):904–9.

66. Purcell S, Neale B, Todd-Brown K, Thomas L, Ferreira MA, Bender D, et al. PLINK: a tool set for whole-genome association and population-based linkage analyses. Am J Hum Genet. 2007;81(3):559–75.

67. O’Reilly PF, Hoggart CJ, Pomyen Y, Calboli FCF, Elliott P, Jarvelin M-R, et al. MultiPhen: Joint Model of Multiple Phenotypes Can Increase Discovery in GWAS. PLOS ONE. 2012;7(5):e34861.

68. DeFries JC, Filipek PA, Fulker DW, Olson RK, Pennington BF, Smith SD, et al. Colorado Learning Disabilities Research Center. Learning Disabilities: AMultidisciplinary Journal. 1997;8:7–19.

69. Willcutt EG, Pennington BF, Olson RK, Chhabildas N, Hulslander J. Neuropsychological analyses of comorbidity between reading disability and attention deficit hyperactivity disorder: in search of the common deficit. Developmental Neuropsychology. 2005;27(1):35–78.

70. Compton DL, Olson RK, DeFries JC, Pennington BF. Comparing the Relationships Among Two Different Versions of Alphanumeric Rapid Automatized Naming and Word Level Reading Skills. Scientific Studies of Reading. 2002;6(4):343–68.

71. Delaneau O, Zagury JF, Marchini J. Improved whole-chromosome phasing for disease and population genetic studies. Nat Methods. 2013;10(1):5–6.

72. Marchini J, Howie B. Genotype imputation for genome-wide association studies. Nat Rev Genet. 2010;11(7):499–511.

73. Perreault L-PL, Legault M-A, Asselin G, Marie-Pierre D. genipe: An automated genomewide imputation pipeline with automatic reporting and statistical tools. Bioinformatics. 2016.

74. Lu Q, Powles RL, Wang Q, He BJ, Zhao H. Integrative Tissue-Specific Functional Annotations in the Human Genome Provide Novel Insights on Many Complex Traits and Improve Signal Prioritization in Genome Wide Association Studies. PLOS Genetics. 2016;12(4):e1005947.

75. Consortium EP. An integrated encyclopedia of DNA elements in the human genome. Nature. 2012;489(7414):57–74.

76. Jernigan TL, Brown TT, Hagler DJ,Jr., Akshoomoff N, Bartsch H, Newman E, et al. The Pediatric Imaging, Neurocognition, and Genetics (PING) Data Repository. Neuroimage. 2016;124(Pt B):1149–54.

77. Alexander DH, Lange K. Enhancements to the ADMIXTURE algorithm for individual ancestry estimation. BMC Bioinformatics. 2011;12:246.

78. Desikan RS, Segonne F, Fischl B, Quinn BT, Dickerson BC, Blacker D, et al. An automated labeling system for subdividing the human cerebral cortex on MRI scans into gyral based regions of interest. Neuroimage. 2006;31(3):968–80.

79. Mehta CM, Gruen JR, Zhang H. A method for integrating neuroimaging into genetic models of learning performance. Genet Epidemiol. 2017;41(1):4–17.

80. Chang CC, Chow CC, Tellier LCAM, Vattikuti S, Purcell SM, Lee JJ. Second-generation PLINK: rising to the challenge of larger and richer datasets. GigaScience. 2015;4(1):7.

